# Agricultural land use and lake morphology explain substantial variation in water quality across Canada

**DOI:** 10.1101/2022.08.29.505280

**Authors:** Joe R. Sánchez Schacht, Paul W. MacKeigan, Zofia E. Taranu, Yannick Huot, Irene Gregory-Eaves

**Affiliations:** Department of Biology, McGill University, Montreal, Quebec, Canada; Aquatic Contaminants Research Division, Environment and Climate Change Canada, Montreal, Quebec, Canada; Département de géomatique appliquée, Université de Sherbrooke, Montreal, Quebec, Canada; Group for Interuniversity Research in Limnology and aquatic environment, Montreal, Quebec, Canada; Department of Natural Resource Sciences, McGill University Macdonald Campus, Sainte-Anne-de-Bellevue, Quebec, Canada

**Keywords:** Agriculture, Canada, eutrophication, lake, urban, water quality.

## Abstract

Despite decades of research and mitigation efforts, declines in freshwater quality resulting from anthropogenic nutrient input remain a persistent issue worldwide. Canada has the greatest number of freshwater lakes in the world, yet we have a limited understanding of the magnitude and scale at which most lakes have been affected by human activities, namely Land Use/Land Cover (LULC) alterations. In response, the NSERC Canadian Lake Pulse Network has compiled the first nationwide systematic database of lake quality metrics by surveying 664 lakes across 12 ecozones over three years. To assess the influence of catchment development on water quality and its spatial variation, we built models quantifying the association between watershed LULC and water quality. We found that agricultural and urban land use explained the greatest proportion of variation in water quality among LULC categories (R^2^ = 0.20–0.29). Overall, our study highlights that drivers of water quality are similar across regions; however, baseline conditions vary, so freshwater ecosystem management strategies must consider their geographic context to better predict where water quality thresholds will be surpassed.

## Introduction

Despite decades of research into the drivers of lake water quality and evolving land management strategies, cultural eutrophication remains one of the leading threats to freshwater ecosystems across the globe (Carpenter et al., 1998; Beusen et al., 2016; Reid et al., 2019). Influxes of limiting nutrients, specifically nitrogen (N) and phosphorus (P), from point sources (i.e., sewage effluent and/or factory discharges) are widely recognized as important contributors to the eutrophication of freshwater ecosystems. Often, the diversion of these point-source nutrients leads to improvements in water quality (Jeppesen et al., 2005; Kronvang et al., 2005). However, mitigation of point sources does not always result in reduced nutrient and algal biomass concentrations in lakes, even in cases where combined (N and P) nutrient loading reduction programs have been applied to lakes and their catchments (Özkundakci et al., 2011; McCrackin et al., 2017). Nonpoint, or diffuse, sources of nutrient inputs remain the main cause of eutrophication worldwide (Reid et al., 2019) partly due to the difficulties in quantifying and regulating them: fluxes can vary temporally (Medalie et al., 2012) and spatially (Lencha et al., 2022) and are often regulated by management strategies with conflicting purposes such as food production and natural resource protection (Wunderlich and Martinez, 2018). Furthermore, annual inputs of N and P from agricultural activities often exceed the removal by harvested crops (MacDonald et al., 2011; Spiess, 2011), leading to a surplus of highly mobile nutrients in soils (GOC, 2016) that can leach into downstream water bodies (Cassman and Dobermann, 2022). The accumulation of soil P and N can result in legacy loading that can take decades to deplete (Grimvall et al., 2000; Van Meter et al., 2016).

LULC in watersheds have previously been linked to variation in lake water quality across the world. More specifically, agriculture and urbanization have been linked to poorer water quality (i.e., eu- or hyper-eutrophic status, increases in Chl-*a*, TP, TN, ions, conductivity, DOC; decrease in Secchi depth) compared to natural landscapes (Taranu and Gregory-Eaves, 2008; Nielsen et al., 2012; Read et al., 2015; de Mello et al., 2020), primarily through higher concentrations of limiting nutrients and their symptomatic cyanobacterial blooms (Doubek et al., 2015; Almanaza et al., 2019). Nevertheless, the relationship between watershed LULC and lake water quality varies significantly over time and space (de Mello et al., 2020) and is often modulated by factors such as precipitation and temperature (Collins et al., 2019), flow regime (Mbonde et al., 2015; Abirhire et al., 2016), lake morphology (i.e., area, depth, altitude; Taranu and Gregory-Eaves, 2008; Read et al., 2015), and bedrock geology (Dillon and Kirchner, 1975; Griffith, 2014). Therefore, studies at a large spatial scale must strive to disentangle the regional and local drivers of water quality. Ultimately, quantifying the effect of nonpoint nutrient loading on water quality across different Canadian ecozones will be instrumental in helping us predict which regions will be more susceptible to lake eutrophication in the event of future LULC modifications.

Our overarching goal is to provide a more robust understanding of how freshwater ecosystems respond to varying levels of anthropogenic stress in a heterogeneous and expansive landscape. By using water physiochemistry and LULC data from 664 Canadian study lakes and watersheds from the NSERC Canadian LakePulse national database (i.e., hereafter referred to as LakePulse), we quantified the spatial variation in water quality (mainly TP, TN, major ion, and Chl-*a* concentrations) among lakes as explained by watershed land use. We tested the hypothesis that the impact of LULC on lake water quality varies across ecozones due to differences in lake and watershed morphology, vegetation, climate, and geology. We applied both linear and nonlinear models to quantify the variation in water quality explained by various LULC categories, including known stressors such as agriculture and urbanization, across ecozones while also considering lake and watershed morphology (altitude, circularity, area, depth, and shoreline slope). Furthermore, we determined thresholds of agricultural and urban land use that were correlated with significantly poorer water quality parameters.

## Materials and methods

### Lake data

Over the span of three summers (June to September) from 2017 to 2019, Lake Pulse teams sampled 664 lakes across all ten Canadian provinces and two territories (Yukon and Northwest Territories). Approximately 600 lakes were in the nine southernmost ecozones (Pacific Maritime, Montane Cordillera, Semi-Arid Plateaux, Prairies, Boreal Plains, Boreal Shield, Mixedwood Plains, Atlantic Maritime, and Atlantic Highland; Figure 1), while the remaining lakes were situated in three northern, and more remote ecozones (Taiga Plains, Boreal Cordillera, and Taiga Cordillera). Target lakes were within 1 km from a road and selected using a random stratified design, controlling for three levels of human impact (low, medium, high; calculated from watershed LULC categories) and three lake sizes (0.1–0.5, 0.5–5, 5–100 km^2^; see Huot et al., 2019). In a few cases, randomly selected lakes were replaced with lakes in the same size and human impact categories because they had associated historical data or because they were important to certain LakePulse partners.

**Figure 1.**
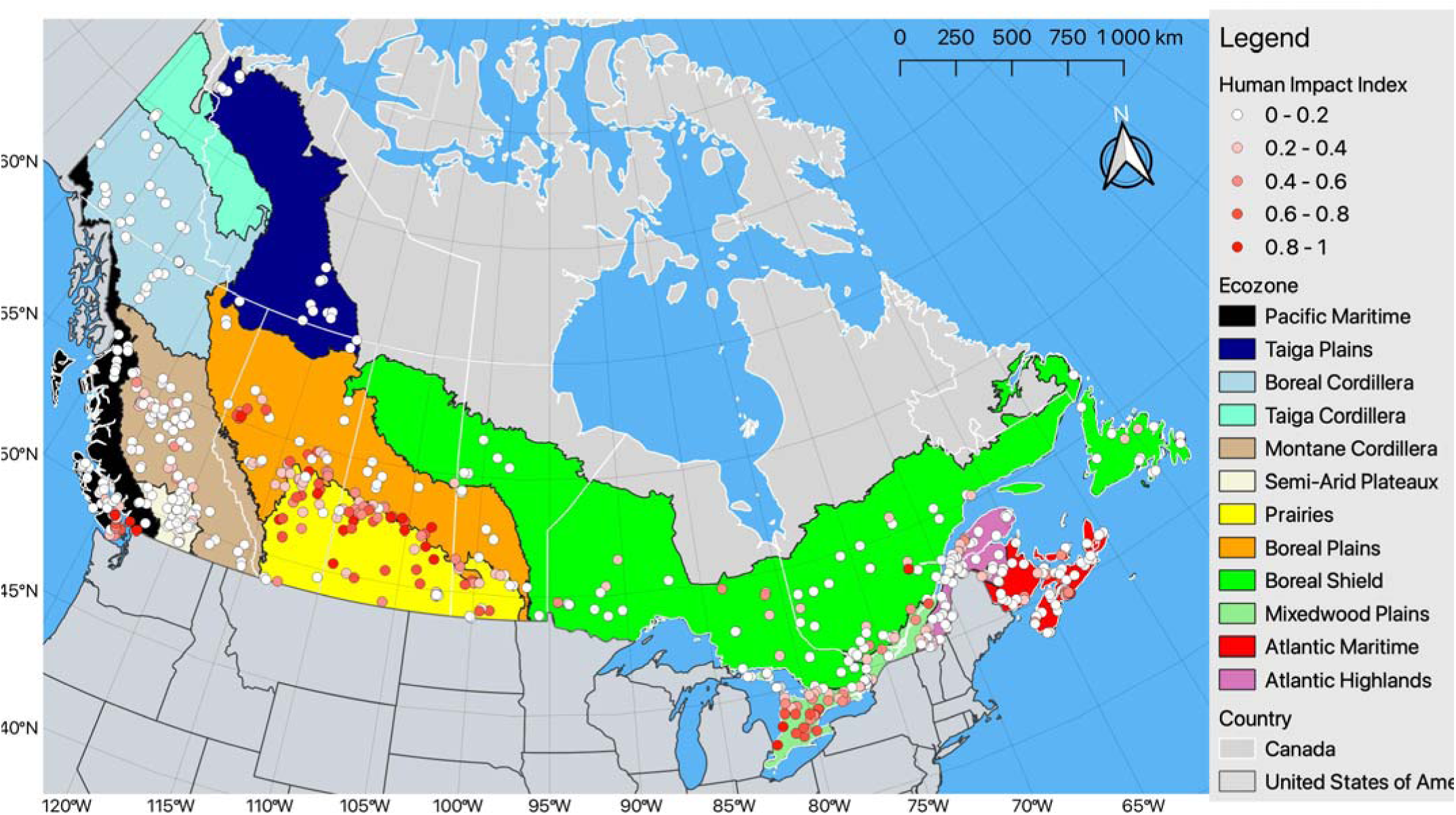
Ecozone map of the 664 sampled watersheds shaded by human impact index. The bulk of sampling was conducted in the nine southernmost main ecozones (Pacific Maritime, Montane Cordillera, Semi-Arid Plateaux, Prairies, Boreal Plains, Boreal Shield, Mixedwood Plains, Atlantic Maritime, and Atlantic Highland), while sampling was limited in the three northernmost and remote ecozones (Taiga Plains, Boreal Cordillera, Taiga Cordillera). This map was illustrated with QGIS 3.0 (QGIS.org, 2022) using shape file and boundary file data from Natural Earth (2022). The projection used is NAD83/Canada Atlas Lambert.

From the deepest point of each lake, survey teams recorded dozens of physical and chemical variables (Table 1). Water quality variables included Secchi depth, dissolved hypolimnetic oxygen, specific conductance, magnesium, potassium, sodium, sulfate, soluble reactive P, total P, total N, Chlorophyll *a*, dissolved organic carbon, and dissolved inorganic carbon. Samples for most water quality variables were taken from the photic zone (2x Secchi depth up to 2 meters), and additional variables physico-chemical (e.g., temperature, specific conductance) were measured across the full water column. In an effort to standardize lake assessment and variable collection protocols across North America, the sampling process was modelled after the NLA in the United States. For full details of the field protocol, see the NSERC Canadian Lake Pulse Network Field Manual (2021). For the purposes of this study, lakes were categorized by their trophic status (oligotrophic, mesotrophic, eutrophic and hypereutrophic) using TP thresholds (Wetzel 2001).

**Table 1.**
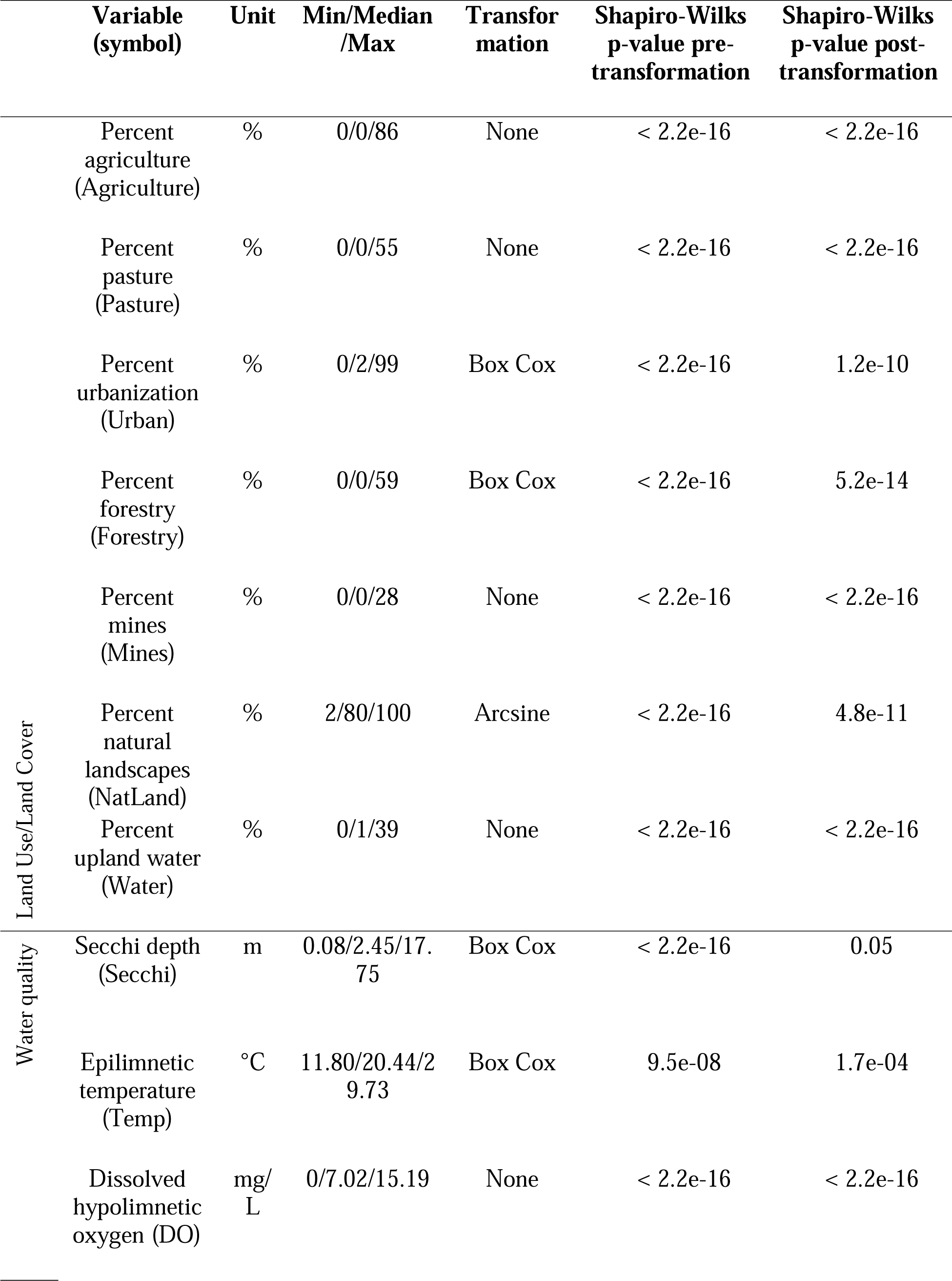
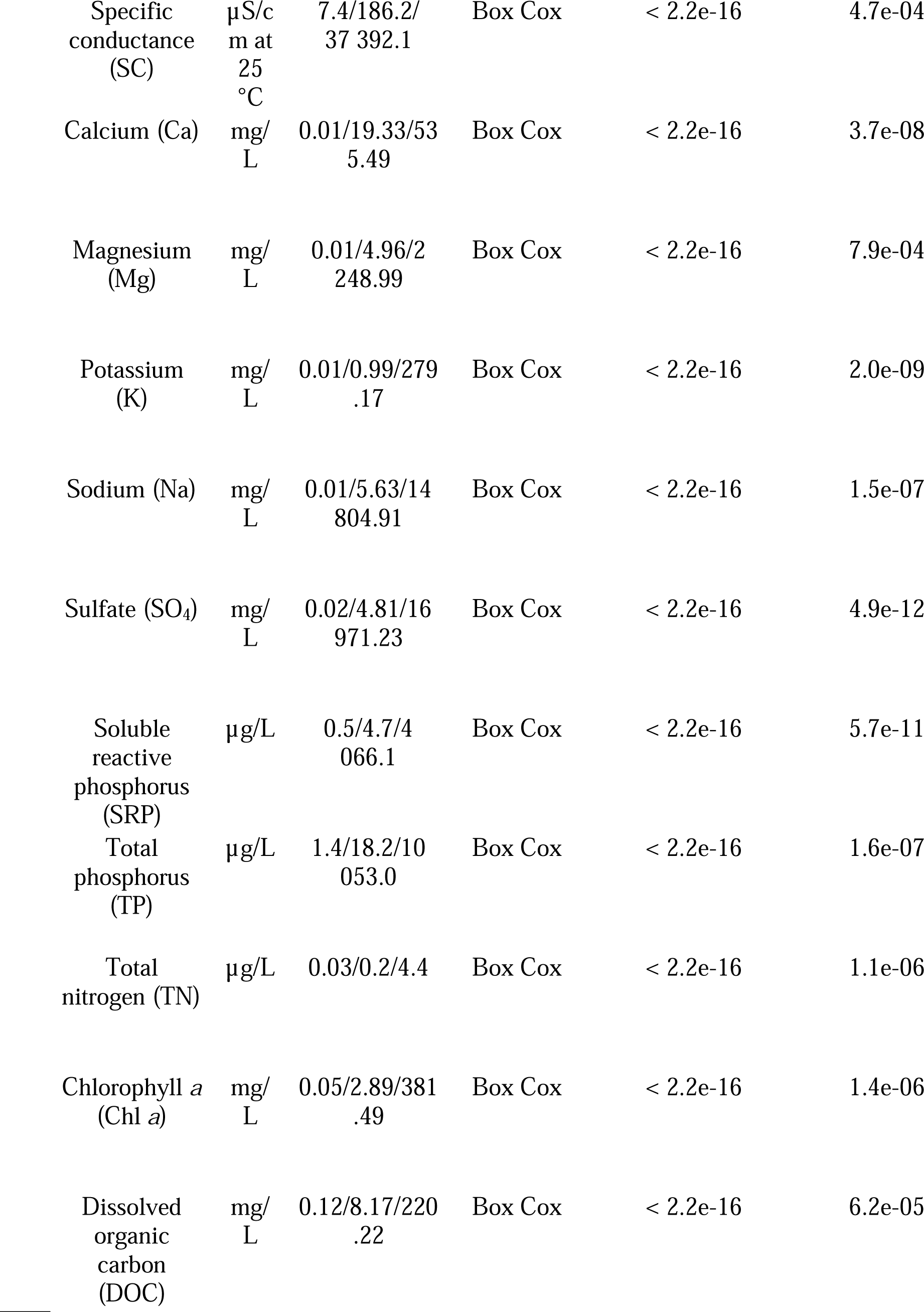
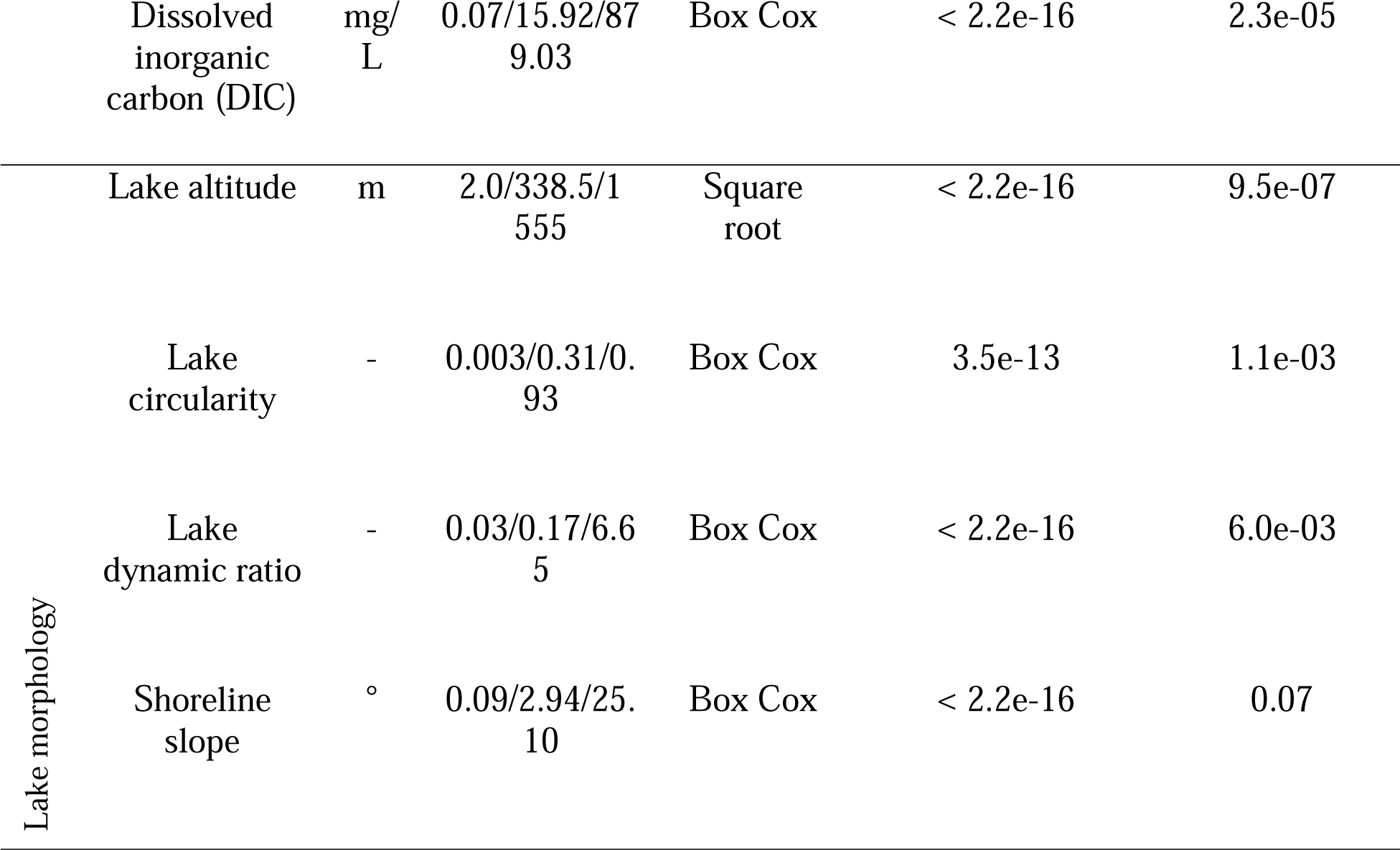
Distribution and transformation coefficients of land use/land cover, water quality, and lake morphology variables for use in the principal component analysis, the redundancy analysis, and the variation partitioning (N = 664 lakes).

### Watershed LULC and morphological data

LULC information for each lake’s watershed was obtained from a variety of sources, including the Annual Space-Based Crop Inventory for Canada 2016 (GOC, 2017) and the Land Use 2010 database (GOC, 2015a), and analyzed using geomatic software (see Huot et al., 2019). Each lake’s respective watershed was delineated to a 30 m^2^ pixel size. Watershed delineation used flow directions calculated with the Canadian Digital elevation Model (GOC, 2015b). LULC type (Agriculture, Pasture, Urban, Forestry, Mines, Natural landscapes, and Upland water) was extracted for each pixel from various public databases and an aggregate watershed human impact was derived based on these LULC data (see Huot et al., 2019 for details). A human impact index was calculated by assigning relative scores to each form of LULC (see Supplemental Table S1) and represents the average value across all pixels in the watershed. Morphometric variables were measured for a subset of lakes and watersheds, four of which were selected *a priori* as covariates in our statistical models: lake altitude, circularity, dynamic ratio (estimated by taking the square root of the lake area:mean depth ratio; Håkanson, 1982; Ferland et al., 2012), and the slope of a 100 m land buffer around the shoreline. Missing data was replaced with the corresponding ecozone median.

### Statistical analyses

All statistical analyses were performed using R v4.0 software (R Development Core Team, 2020). Since several of our analyses assumed a normal data distribution, we examined the distribution of each response variable using the Shapiro-Wilks test combined with a visual inspection of histograms. For all variables that did not meet the test’s normality assumption (p > 0.05), we applied a Box Cox (Borcard et al., 2018), logarithmic, or square root transformation. The Box Cox transformations were computed with the R package *geoR* (Ribeiro et al., 2020).

The transformed variables were then evaluated again with the Shapiro-Wilks test and a histogram inspection, and the transformation that was closest to being normal was kept (specific transformations, where appropriate, are included in Table 1). To improve model fit, the same test for normality and transformation exercise were applied to the LULC variables, for which we additionally attempted an arcsine transformation. For the principal components, redundancy, and variation partitioning analyses, LULC variables with zero values were offset by adding 0.00001 when Box Cox or logarithmic transformations were applied. However, the raw LULC values were used for the multivariate regression tree (MRT), cascade multivariate regression tree (cMRT), and generalized additive models (GAMs). We found no significant collinearity (>0.7 Pearson correlation or >10 VIF) among the predictors. However, watershed proportions of natural landscapes and mines were omitted as predictors to prevent redundancy and convergence issues in our models.

To visualize the correlations among lake water quality variables, and proximities in ordination space among ecozones, we conducted a principal component analysis (PCA) using the *rda* function of the R package *vegan* (Oksanen et al., 2019). All response variables were transformed (Table 1), then standardized (scaled to zero mean and unit variance) prior to performing the PCA. LULC variables were fitted passively on the PCA to illustrate their association with water quality parameters. We used the broken-stick method (Jackson, 1993) to determine the PC axes that accounted for a significant portion of the variation observed. Since most of the variation was explained by PC axis 1, we applied an analysis of variance (ANOVA) approach to quantify the observed variability in axis scores between ecozones and post-hoc Tukey’s tests to determine significant differences.

With the goal of exploring the relationships between different LULC and water quality parameters, we then performed a multivariate canonical ordination (i.e., a redundancy analysis [RDA]) using the *rda* function of the R package *vegan* (Oksanen et al., 2019). We set LULC proportions as predictors and water quality measurements as response variables. RDAs assume linear relationships between explanatory and response variables. Therefore, the RDA was performed using the transformed variables (Table 1). All response variables were then standardized (scaled to zero mean and unit variance) prior to performing RDA and significant predictors were selected using forward selection (Legendre and Legendre, 2012). The omitted LULC variables (Natural landscapes and Mines) were also fitted passively on the RDA to illustrate their association with water quality parameters. Since most of the variation was explained by RDA axis 1, we used the same ANOVA and Tukey’s tests approach as for the PCA to quantify variation among ecozones.

To disentangle the added influence of lake and watershed morphology on these relationships, we performed a variation partitioning using the *varpart* function of the R package *vegan* (Oksanen et al., 2019). Here, we considered the unique vs. combined proportion of variation in lake water quality that LULC (all categories) and morphology explained, where altitude, lake circularity, shoreline slope, and the dynamic ratio were bundled into the morphology metric.

Finally, to examine potential nonlinear relationships that were not captured by the linear RDA, we first investigated the direct relationships between the raw proportion of agriculture in the watershed and three commonly used indicators of water quality (Chl-*a*, TP, and TN; all log transformed for interpretability) using generalized additive models (GAMs). We chose GAMs due to the hypothesized non-linearity of these relationships in ecozones with either low or high human impact. We also performed a multivariate regression tree (MRT) using the R package *mvpart* (Therneau and Atkinson, 2014) to test for threshold responses (Ouelette and Legendre, 2013) using all response variables (transformed and standardized environmental data). We used the cross-validation relative error (CVRE), the ratio of the variation unexplained by the model to the total variation in the response, to select the most parsimonious tree (i.e., the tree with the least number of nodes whose CVRE value was within one standard error of the tree with the lowest CVRE; De’Ath, 2009). As an alternative way to calculate specific threshold LULC proportions, we also applied a cascade MRT (Ouellette et al., 2012), which permitted us to impose a nested structure on the analysis and partition the explanatory power of two independent datasets, a 664 × 7 matrix of raw LULC percentages and a 664 × 1 ecozone vector, to identify regional variability in the relationship between LULC and water quality (see Supplemental Figure S5). Essentially, the model computes trees in the following order: (1) it initially separates all sites according to ecozone, highlighting significant geographic discrepancies in lake water quality; and (2) it then, for each branch (i.e., group of ecozones), uses the environmental data to further separate sites with respect to LULC proportions.

## Results

### Local to regional variation in water quality, lake morphology, and LULC

We found significant variation in lake morphology and LULC among ecozones. For example, the proportion of agriculture in a given lake’s watershed was greatest in the Prairie, Boreal Plains and MixedWood Plains ecozones (Supplemental Figure S1). Likewise, we detected substantial differences in altitude among ecozones (Supplemental Figure S2). Given this heterogeneity in environmental conditions across the country, it is not surprising that we also found substantial variation in water quality metrics (Supplemental Figure S3).

To summarize the variation in lake water chemistry data, we conducted a PCA and found that PC axis 1 (PC1) explained 51.2% of the variance. Ions (Na^+^, K^+^, Ca^2+^, Mg^2+^, and SO ^2−^), specific conductance (SC), and dissolved inorganic carbon (DIC) had large loadings on PC1 and were associated with watersheds with more urban development and agriculture (Figure 2a). TN, TP, DOC, soluble reactive phosphorus (SRP), and Chl-*a* also loaded strongly on PC1, but were associated with higher proportions of agriculture and pasture (Figure 2a). PC2 distinguished sites with a higher proportion of natural landscapes that had lower TP, TN, DOC and DIC but greater Secchi disk depths (Figure 2a). Sites in the Prairies and Boreal Plains (negative PC1 loadings), tended to differ from sites in the Mixedwood Plains or the Atlantic and Pacific Maritimes (tendency for positive PC1 loadings; Figure 2b). This difference was supported by a one-way ANOVA which showed that lakes within the Prairies and Boreal Plains ecozones had significantly different PC1 loadings than all other ecozones, distinguishing them from lakes across the rest of Canada (F_(11, 652)_ = 103.5, p < 0.001; Tukey HSD: p < 0.05; Figure 2c).

**Figure 2.**
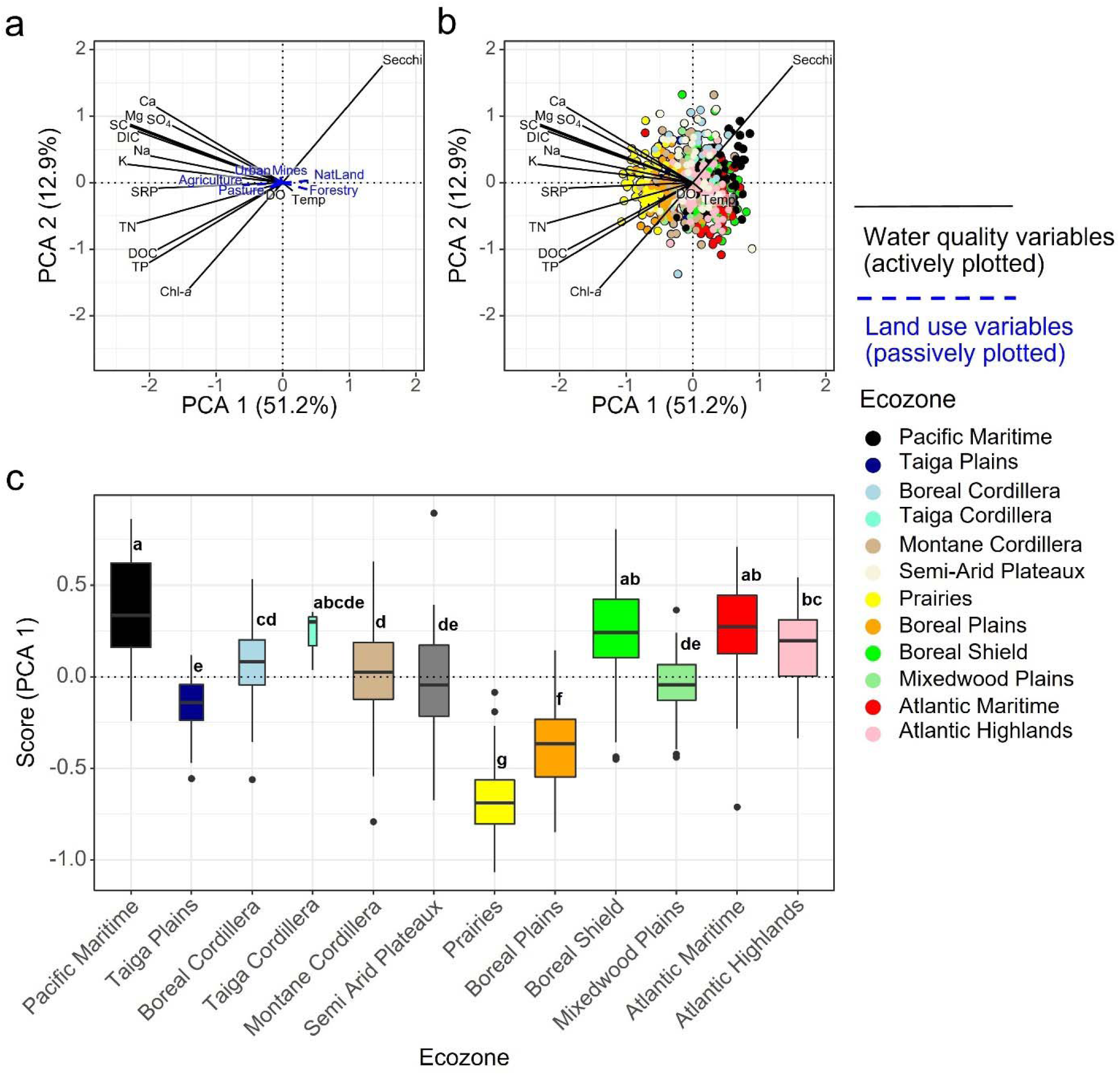
Principal component analysis (PCA) of lake variables from the LakePulse dataset (N = 664). The PCA is plotted twice to highlight the distribution of a) vectors of LULC types plotted passively (coloured in blue, dashed lines) and b) site scores, colour-coded according to ecozone. c) Boxplot of site scores from PC axis 1 grouped and colour-coded by ecozone, where the width of the boxes reflects sample size ranging from 3 to 91 lakes. Letter superscripts denote significantly different groups determined via a post-hoc Tukey’s test.

### Agriculture and pasture explain the most variation in water quality

The redundancy analyses with LULC proportions as the explanatory variables revealed similar gradients as the PCA, with an important difference: the resulting vectors for agriculture and urbanization were orthogonal in the RDA. The analysis ordinated the various sites into three significant axes that explain 23.7% of the variance. Axes 1, 2 and 3 explained 21.1%, 1.9% and 0.7% of the variance, respectively. The proportion of agriculture or pasture were found to have strong loadings on RDA axis 1, and were correlated with TN, TP, DIC, DOC, SRP, Chl-*a*, all ion concentrations, and hypolimnetic oxygen (DO), whereas Secchi disk depth had a negative RDA axis 1 loading and was associated with forestry activity and natural landscapes (Figure 3a). In contrast, urbanization was positively associated with RDA axis 2 and correlated with higher epilimnion temperatures and sodium, but lower hypolimnetic oxygen concentrations (Figure 3a). The one-way ANOVA of RDA 1 scores revealed that sites in the Prairies and Boreal Plains, strongly associated with agriculture and pasture, differed significantly from other ecozones (F_(11, 652)_ = 99.4, p < 0.001; Tukey HSD: p < 0.05; Figure 3b).

**Figure 3.**
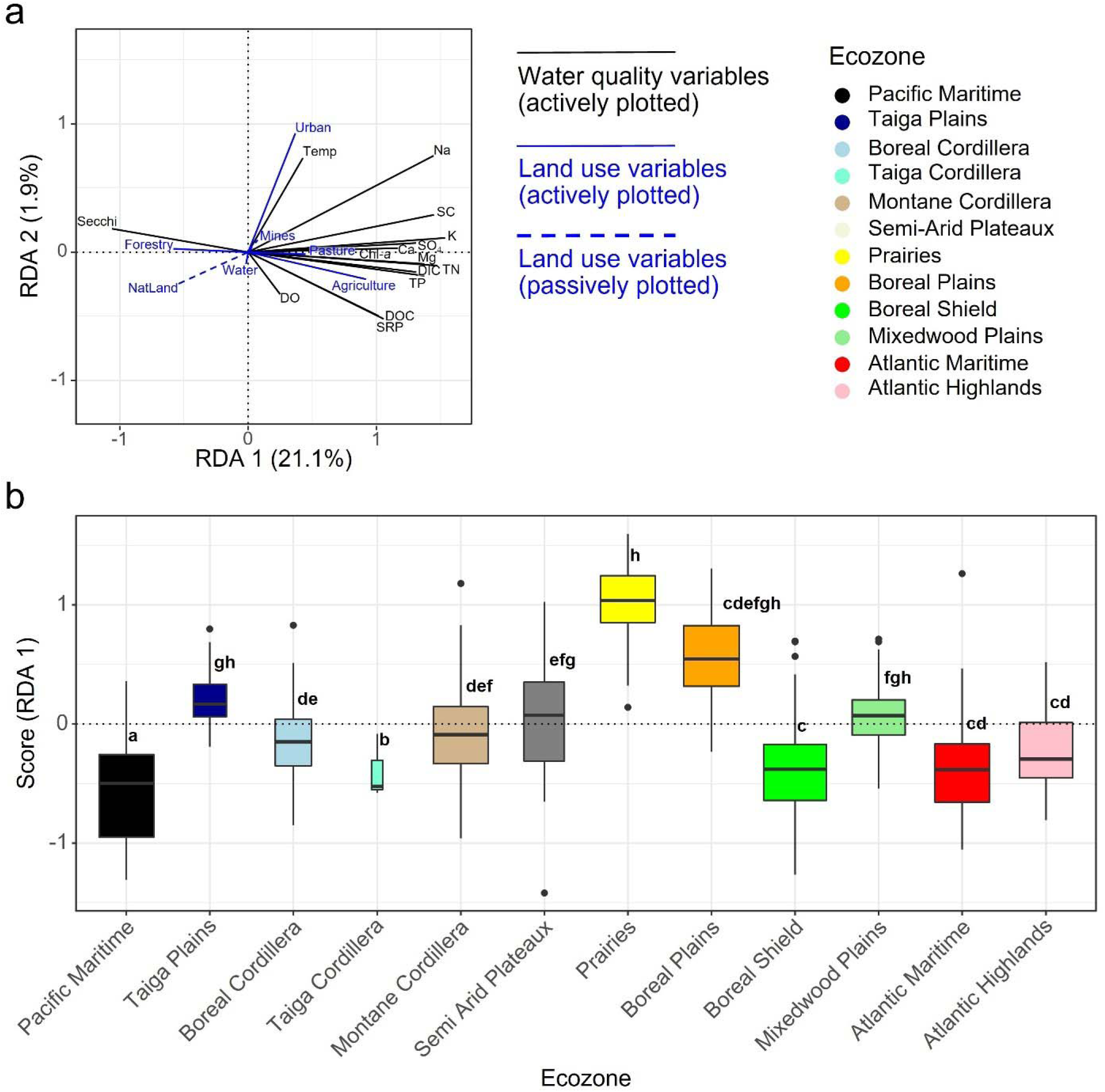
Redundancy analysis examining the relationships between water quality variables (response variables) and LULC metrics (predictors; N = 664). Response variables were transformed and standardized prior to analysis. The analyses are divided into two plots to highlight the distribution of a) vectors of LULC variables (coloured in blue), and b) boxplot of site scores for RDA axis 1 grouped by ecozone, where the width of the boxes reflects sample size ranging from 3 to 91 lakes. Letter superscripts denote significantly different groups determined via a post-hoc Tukey’s test.

The variation partitioning including altitude, lake circularity, shoreline slope, and the dynamic ratio as covariables describing lake and watershed morphology further emphasized the importance of LULC as a predictor of lake water quality. LULC uniquely explained a total 15% of the variation in water quality, morphology uniquely explained 11%, while 9% of the variation was co-explained by both groups of variables (Figure S4).

### Variable thresholds and plateau responses in the Canadian Prairies

GAMs showed that agricultural land use had varying effects on lake water quality indicators (Chl-*a*, TP, and TN concentrations) depending on ecozone (i.e., local bedrock geology, lake and watershed morphology). Mean values for TP and TN in watersheds with little to no agriculture (intercepts) were relatively higher for the Prairies than for the Mixedwood Plains and Boreal Shield ecozones (Figure 4), and the response to greater proportions of agriculture in the watershed tended to plateau in the Prairies and Boreal Plains relative to trends observed in other ecozones.

**Figure 4.**
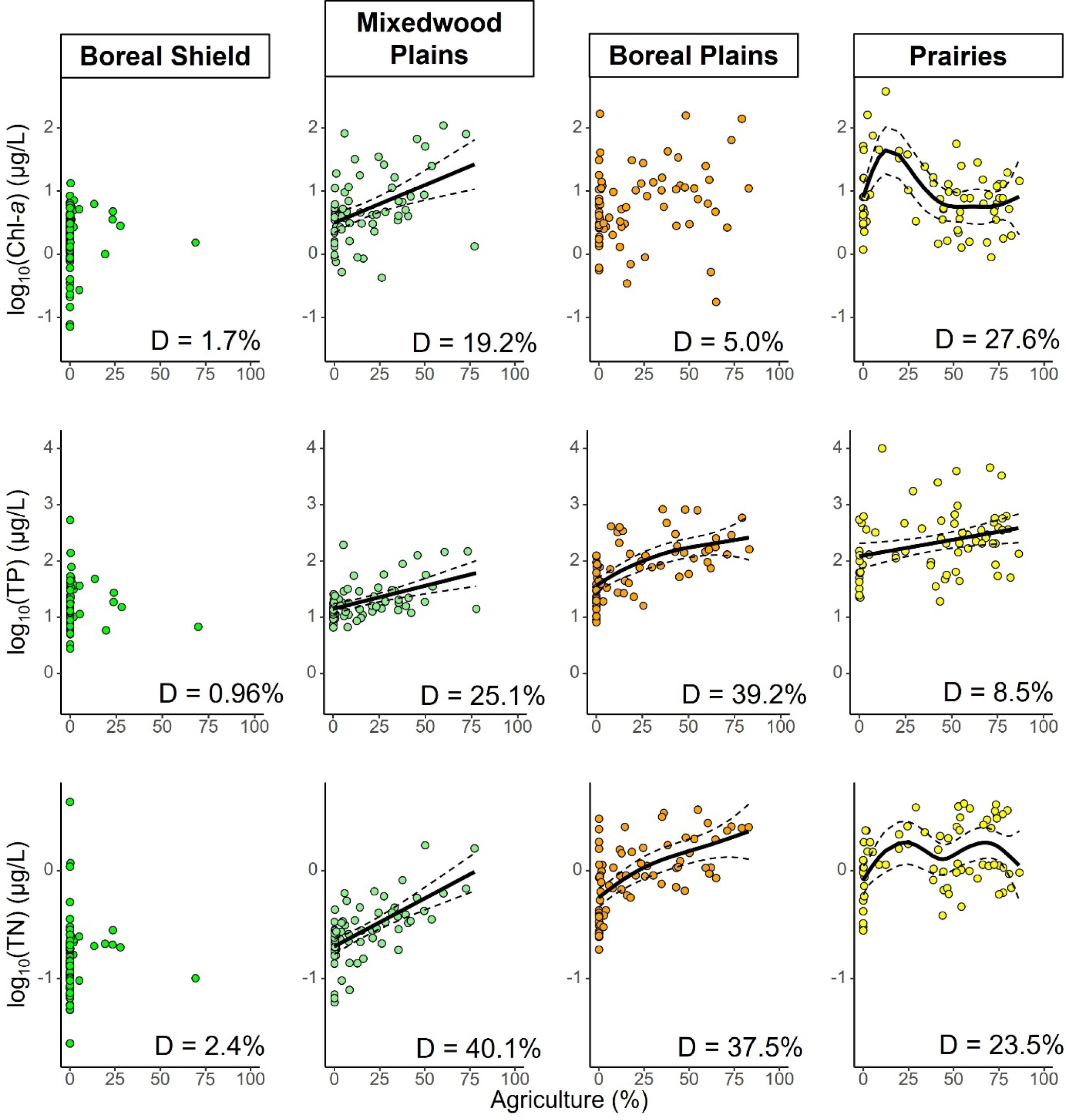
Generalized additive models of the relationships between % Agriculture and total phosphorous (TP), total nitrogen (TP), and chlorophyll a (Chl-*a*) concentration in four populous ecozones along an increasing human impact gradient. Dashed lines represent the 95% confidence intervals. Sites are coloured according to ecozone. Deviation explained by each model (D) is shown at the bottom right of each figure.

The multivariate regression tree analysis identified threshold values for agriculture and urbanization where we could expect pronounced changes in lake water quality across the country (Figure 5). In particular, the presence of agriculture (>0.1% of the watershed) was identified as the most important predictor of water quality, whereby sites with minimal agriculture and urbanization had higher than average (i.e., the baseline within each branch of the MRT) Secchi depths, lower than average concentrations of ions, nutrients, and lower epilimnetic temperatures (Figure 5). Among sites with no to very little agriculture, lake watersheds with some urban land use (i.e., >1.3%) had ions, nutrients and water temperatures that were closer to the national average (Figure 5). Water quality in sites with moderate agriculture (i.e., between 0.2% and 36%) was similar to that of sites with some urban land use, albeit with slightly higher than average values (Figure 5). Sites with substantial agriculture (i.e., >36%) had significantly higher than average ion and nutrient concentrations while Secchi depths were considerably lower than average (Figure 5).

**Figure 5.**
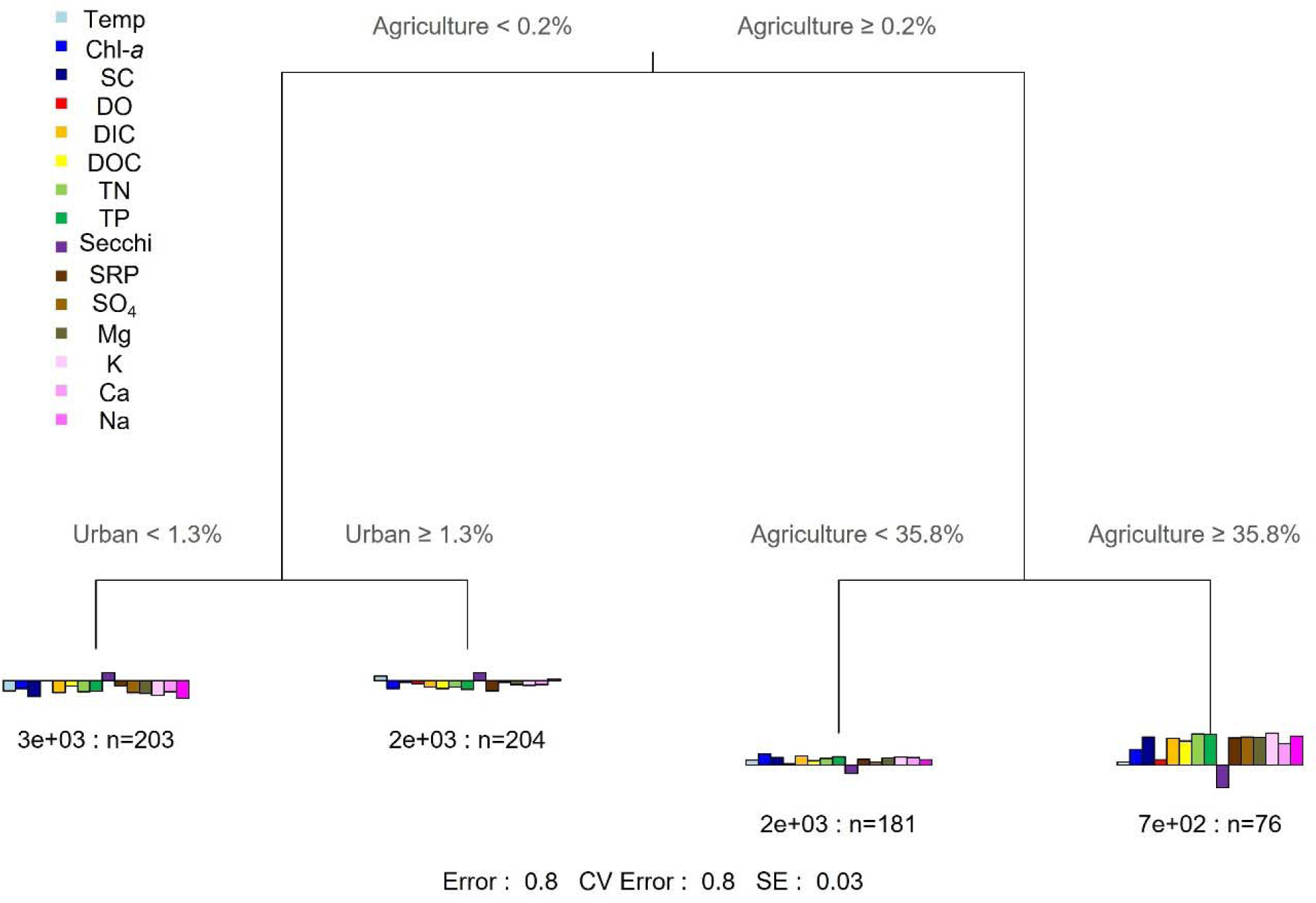
Multivariate regression tree of the transformed and standardized water quality variables, constrained by the LULC variables (R^2^ = 0.20; N = 664).

The cascade regression tree (cMRT), which was performed to consider potential regional differences, clearly identified the difference in lakes of the Prairie ecozone from other regions. The cMRT also increased the proportion of variation explained slightly relative to the MRT (R^2^ = 0.29; Figure S5), but identified similar thresholds in land use proportions. In the Prairies, water quality varied slightly by the presence of agriculture in the watershed, since catchments with even modest agricultural land (i.e., >0.2%) had ion and nutrient concentrations above the national average (Figure S5). In other ecozones, water quality parameters only reached above average values when the proportion of agricultural land surpassed 0.8%, although lakes in watersheds with proportions between 0.8% and 29.7% remained relatively close to the national average. Additionally, water quality in catchments with less than 0.8% agriculture varied by their proportion of urbanization, with sites over 6.6% urban land showing higher epilimnetic temperatures and specific conductance, but lower hypolimnetic oxygen compared to the national average as well as their less urbanized counterparts (Figure S5).

## Discussion

Our analyses of 664 lakes from twelve ecozones, spread across a total study area of 412 052 km^2^, revealed considerable variation in lake water quality parameters across the nation’s ecozones, with the Canadian Prairies showing significantly different lake water quality compared to the rest of Canada. The higher proportions of agriculture and pasture in the Prairies suggest an important link (Figures 2 and 3), although it is also clear that other factors are at play given the heightened baseline nutrient values for this ecozone (Figure 4). Both linear and nonlinear multivariate regression methods identified agriculture and urbanization as the main land use drivers of lake water quality across a wide geographic gradient (Figures 3, 5 and S2). Variation partitioning with lake and watershed morphology factors as covariates demonstrated that this relationship was modulated by geological and topographical factors, some of which were inherently correlated with LULC (Figure S4). Multivariate regression tree models also identified the thresholds at which agricultural and urban land use are associated with poorer water quality, and highlighted differences between the Prairies and other ecozones (Figures 5 and S2). Furthermore, we found that nonlinear models explain more variation in water quality than linear models, which underscores the importance of further examining ecological causes for nonlinearity in the relationship between LULC and water quality metrics.

### LULC, morphology and lake water quality vary among Canadian ecozones

We observed that most lakes in the Prairies, and at least some in the Boreal Plains, had significantly different water quality values relative to the rest of the country. Specifically, lakes in the Prairies and Boreal Plains had higher than average concentrations of TP, TN, major ions, dissolved carbon and algal biomass (inferred from Chl-*a*), and lower than average Secchi depths and hypolimnetic oxygen, all of which contributed to substantial variation in the national dataset (Figures 2b and 2c). These marked discrepancies in water quality for the Prairies and Boreal Plains ecozones are likely the result of a higher local prevalence of agriculture and a difference in baseline nutrient conditions (Prairies TP_median_ = 204.1 µg/L, Canada TP_median_ = 18.2 µg/L; Figures 4 and S3; Taranu and Gregory-Eaves 2008). There are multiple factors that might contribute to this regional discrepancy in lake responses to LULC development. For one, the region’s flat catchment topography (Last and Ginn, 2005; Bharath and Elshorbagy, 2018), naturally nutrient-rich geology (Taranu and Gregory-Eaves, 2008; Griffiths et al., 2022) and its changing hydrology (i.e., declining river flows due to a reduction in yearly persistence and maximum depth of local snowpacks; Schindler and Donahue, 2006; see Ireson et al., 2015 for a general overview of the region’s shifting hydrology) may further modulate the effect of agricultural development and urban expansion (Leavitt et al., 2006; Spence et al., 2019). Indeed, the combined effects of increasing agricultural and urban development and the Prairie region’s flat, nutrient-rich landscape on lake water quality were supported by our variation partitioning analysis, where LULC explained a considerable amount of the variation (15%) in water quality alongside lake and watershed morphology (11%), and their interaction (9%; Figure S1). There is also a well-known tendency for eutrophication to occur in shallow lakes set in flat catchments, and this, coupled with the natural productivity of the Prairies, likely modulated the relationship between LULC and lake water quality. Consistent with these findings, we found that TP, TN, and DOC were higher than the national average, and that DO was lower than average in the LakePulse Prairie sites (Figure S3). In addition, climatic variability (e.g., Chinook winds locally and droughts regionally; see Lemmen et al., 1997) has been shown to profoundly affect continental lakes (Schindler and Donahue, 2006), leading to variations in water quality across different spatial and temporal scales (Pham et al., 2009). However, Vogt et al. (2018) have also shown that internal lake characteristics (i.e., limiting nutrients, oxygen content, DOC) can have an even larger influence on the water quality of lakes in this region relative to that of climate and meteorology.

There is a growing body of evidence that has identified that long-term anthropogenic impacts may have disproportionately altered the productivity and water physiochemistry of Prairie lakes relative to other ecozones, leading to the present-day dichotomy in baseline nutrient levels and water quality. For example, time-series analyses revealed widespread increases in algal biomass since pre-industrial times (Taranu et al., 2015; Ho et al., 2019; Griffiths et al., 2022), and indicate that in highly developed and economically important ecozones, such as the Prairies and the Mixedwood Plains, these trends have accentuated. Furthermore, analyses of bioindicators preserved in lake sediment records have shown that the highest degree of taxonomic turnover since pre-industrial conditions have been localized in ecozones hosting intense land uses, with agriculture and urbanization being key stressors to those lakes (Griffiths et al., 2021; Griffiths et al., 2022).

### Land use thresholds for lake water quality differ among ecozones

With nonlinear modelling methods, we identified land use thresholds that lead to significant changes in water quality. The initial branch of both the MRT and the cMRT separated sites according to whether agriculture was present in the watershed, suggesting that agricultural land use was the main determinant of water quality across all sites. Interestingly, nested analyses suggested that subsequent inflection points differed between the Prairies sites and the rest of Canada. The MRT and cMRT both identified an agricultural threshold of 30–35% after which a large decrease in water quality was observed (i.e., increase in limiting nutrients, ions, carbon, and algal biomass; Figures 5 and S5). Water quality across all sites was poorer when even a minute proportion of the catchment was devoted to agricultural activities (0.2% in the Prairies, 0.8% in the rest of Canada; Figure S5). However, the differences in water quality were much more pronounced after surpassing the 29.7% agriculture threshold across Canada, relative to exceeding the 0.2% threshold in the Prairies ecozone (Figure S5), suggesting a lessened effect of additional nutrient input on algal biomass in eutrophic and hypereutrophic Prairie lakes. Many lakes in this region were historically productive, being at least mesotrophic (Lemmen et al., 1997; Blais et al. 2000), which might partially explain why the change in water quality is modest in the Prairies, where baseline nutrients are already high.

We also identified a national urbanization threshold of 1–7%, after which we observed altered water quality evidenced by increases in sodium concentrations and epilimnion temperatures (Figures 5 and S5). Since roads were classified as urban land use (Supplemental Table S1), the observed increases in sodium concentrations might be due to various factors such as the widespread practice of road salt application, leaky septic fields, deteriorating infrastructure (Gagnon et al., 2008; CIRC, 2019), or a combination of these factors. Increases in epilimnetic temperatures are consistent with past studies on examining the effect of urbanization on lake water temperature (LeBlanc et al., 1997; Yang et al., 2020), although climate change is also impacting lake temperatures globally (O’Reilly et al., 2015; Woolway et al., 2020). Additionally, the association between urbanization and epiliminetic temperatures could be related to geography as Canadian municipalities are concentrated in the south (and we observed a significant negative correlation between latitude and percent urban land use in our dataset).

Across the body of literature that quantified land use threshold values in association with lake water quality, there is substantial heterogeneity. For example, Qiu and Turner (2015) found an agricultural land cover threshold of 13% using data from a Wisconsin watershed, whereas Baker (2006) reported an evident reduction in water quality at a threshold of 5% urban land use when using a global array of studies. It is possible that the climatic and geomorphological context modifies the influence of land use globally and could help reconcile these seemingly disparate findings, as we have seen in our Canadian dataset. Further refinement of these models could be performed with a closer examination of the type of agriculture (e.g., traditional vs. organic farming, type of crop) and urbanization across different watersheds, to help inform policy development and to provide more data and incentives for farmers.

### Conclusions

Through the application of regression models to an expansive dataset of limnology and LULC proportions, we have shown that while water physiochemistry varies across Canadian lakes, watershed land use has a direct and significant effect on water quality, especially agriculture. We also found that lake water quality responds differently to land use across regions, which can be explained by geology, morphology (with altitude showing the largest variation between ecozones), land use history, and climate. Nonetheless, with further expansion in the amount of land converted to agricultural activities due to population growth and increasing uncertainty with regards to global food security, we predict future declines in lake water quality. Looking forward, initiatives such as high precision, organic, and/or vertical farming might pave the way for more space-efficient agriculture that results in less run-off of nutrients to downstream water bodies (Viana et al., 2022). Since organic farms incur a considerable risk of soil P depletion (Carr et al., 2019), recycling anthropogenic P waste is also being considered as a possible alternative source of nutrients (Nicksy and Entz, 2021).

Nonetheless, agricultural production is an important contributor to Canada’s economy and a critical part of society. For example, the nation’s agricultural sector generated 39.8 billion Canadian dollars in 2020 (GOC, 2021). Given the economic importance of its domestic agricultural production, the federal, provincial, and territorial governments formed the Canadian Agricultural Partnership, investing three billion dollars into federal programs and strategic initiatives over five years (2018–2023) to help strengthen the industry, of which 23% is dedicated to research and innovation (GOC, 2019). Other investments by the Canadian government over the next 10 years are targeted for the adoption, testing, and monitoring of best management practices (BMPs; e.g., no-till, cover crops) that directly or indirectly improve inland water quality (Faust et al., 2018). Yet, voluntary participation in unsubsidized BMPs remains low (Liu and Brouwer, 2022), with Canadian agricultural producers citing costs as an important factor when deciding to implement BMPs (Hassanzadeh et al., 2019). Failing to adopt BMPs and other progressive agricultural strategies can lead to the development and persistence of harmful algal blooms across North America (Munawar and Fitzpatrick, 2018; Graham et al., 2020).

Stakeholder engagement remains crucial to successful water quality management, as the rates of BMP adoption by producers can influence the effectiveness of said practices at a watershed scale (Zammali et al., 2021). Land management strategies must take these particularities into account if we are to preserve crucial ecosystem services such as food production and potable water access worldwide and beyond the foreseeable future.

## Acknowledgements

The authors thank the LakePulse data collection and data curating teams for their provision of the data, as well as Maxime Fradette, Katherine Griffiths, Alexandre Baud, Michelle Gros, Candice Aulard, David Zilkey, Rebecca Garner, and all the Gregory-Eaves lab for their continued support and insightful advice.

## Author contribution statement

J.R.S.S. – Funding acquisition, methodology, formal analysis, visualization, writing – original draft.

P.M. – Investigation, data curation, methodology, validation, supervision, writing – review & editing.

Z.E.T. – Methodology, supervision, writing – review & editing.

Y.H. – Funding acquisition, conceptualization, methodology, project administration, supervision, writing – review & editing.

I.G.-E. – Funding acquisition, conceptualization, methodology, project administration, supervision, writing – review & editing.

## Data availability statement

Data generated or analyzed during this study will be available at https://lakepulse.ca/.

## Table

## Figure captions

## Appendix A: Supplemental Table

**Supplemental Table S1.** Annual Space-Based Crop Inventory for Canada 2016 and Land Use 2010 database categories and corresponding simplified categories of the various LULC types. Simplified categories were used in this article’s analyses.

| Definition | Simplified category |
| --- | --- |
| No Data | NA |
| Water | Water |
| Exposed Land / Barren / Rock / Snow / Ice | Natural Landscape |
| Natural landscape (unclassified) | Natural Landscape |
| Bryoids | Natural Landscape |
| Shrubland | Natural Landscape |
| Herb / Grassland | Natural Landscape |
| Coniferous forest | Natural Landscape |
| Broadleaf forest | Natural Landscape |
| Mixedwood forest | Natural Landscape |
| Forest (undifferentiated) | Natural Landscape |
| Wetland | Natural Landscape |
| Forest Loss (2012-2017) | Forestry |
| Urban / Developed | Urban |
| Sod | Urban |
| Greenhouses | Agriculture |
| Agriculture (undifferentiated) | Agriculture |
| Pasture / Forages | Pasture |
| Too Wet to be Seeded | Agriculture |
| Fallow | Agriculture |
| Cereals | Agriculture |
| Barley | Agriculture |
| Other Grains | Agriculture |
| Millet | Agriculture |
| Oats | Agriculture |
| Rye | Agriculture |
| Spelt | Agriculture |
| Triticale | Agriculture |
| Wheat | Agriculture |
| Switchgrass | Agriculture |
| Sorghum | Agriculture |
| Winter Wheat | Agriculture |
| Spring Wheat | Agriculture |
| Corn | Agriculture |
| Tobacco | Agriculture |
| Ginseng | Agriculture |
| Oilseeds | Agriculture |
| Borage | Agriculture |
| Camelina | Agriculture |
| Canola / Rapeseed | Agriculture |
| Flaxseed | Agriculture |
| Mustard | Agriculture |
| Safflower | Agriculture |
| Sunflower | Agriculture |
| Soybeans | Agriculture |
| Pulses | Agriculture |
| Peas | Agriculture |
| Beans | Agriculture |
| Lentils | Agriculture |
| Vegetables | Agriculture |
| Tomatoes | Agriculture |
| Potatoes | Agriculture |
| Sugar beets | Agriculture |
| Other Vegetables | Agriculture |
| Fruits | Agriculture |
| Berries | Agriculture |
| Blueberry | Agriculture |
| Cranberry | Agriculture |
| Other Berry | Agriculture |
| Orchards | Agriculture |
| Other Fruits | Agriculture |
| Vineyards | Agriculture |
| Hops | Agriculture |
| Herbs | Agriculture |
| Nursery | Agriculture |
| Buckwheat | Agriculture |
| Canary seed | Agriculture |
| Hemp | Agriculture |
| Vetch | Agriculture |
| Other Crops | Agriculture |
| Mines/excavation | Mines |

## Appendix B: Supplemental Figure captions

Supplemental Figure S1. Boxplot distributions of Land Use/Land Cover variables used in this study, per ecozone (N = 664). Boxplots are colour-coded by ecozone. When variables were transformed, y-axis displays transformed units in black and original units in grey.

Supplemental Figure S2. Boxplot distributions of morphological variables used in this study, per ecozone (N = 664). Boxplots are colour-coded by ecozone. When variables were transformed, y- axis displays transformed units in black and original units in grey.

Supplemental Figure S3. Boxplot distributions of water quality variables used in this study, per ecozone (N = 664). Boxplots are colour-coded by ecozone. When variables were transformed, y- axis displays transformed units in black and original units in grey.

Supplemental Figure S4. Variation partitioning of water quality variables (response variables) among LULC proportions and lake morphology (explanatory variable groups; R^2^ = 0.35; N = 664). Water quality variables were transformed and standardized prior to analysis. Percentages inside circles represent the total variation explained by the explanatory variables included in each group, with each group being distinguished by its own colour.

Supplemental Figure S5. Cascade multivariate regression tree of the transformed and standardized water quality variables, constrained by the LULC proportions (R^2^ = 0.29; N = 664). The model first splits the dataset geographically, then by LULC in response to water quality differences.

## Appendix C: Supplemental Figures

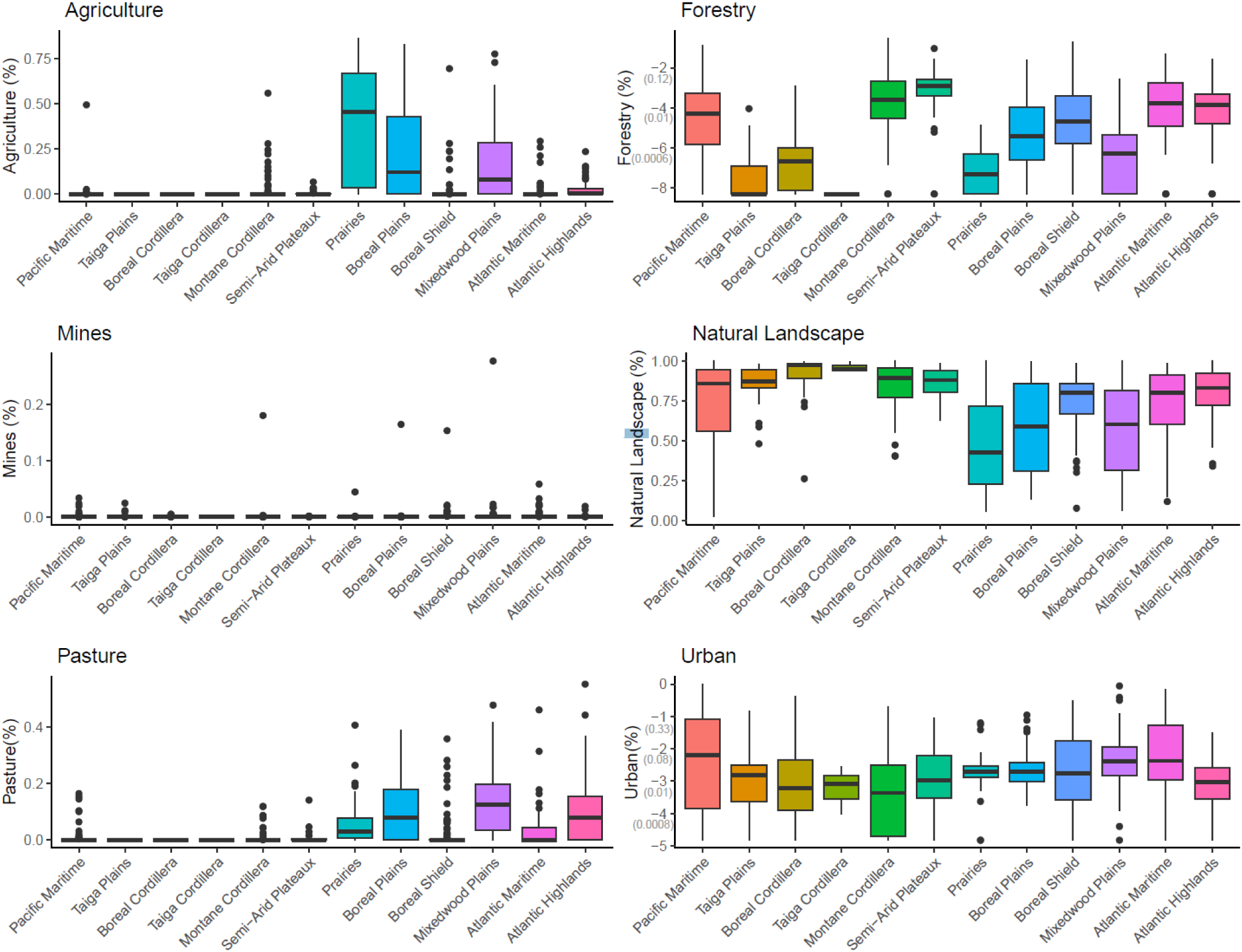
Supplemental Figure S1.

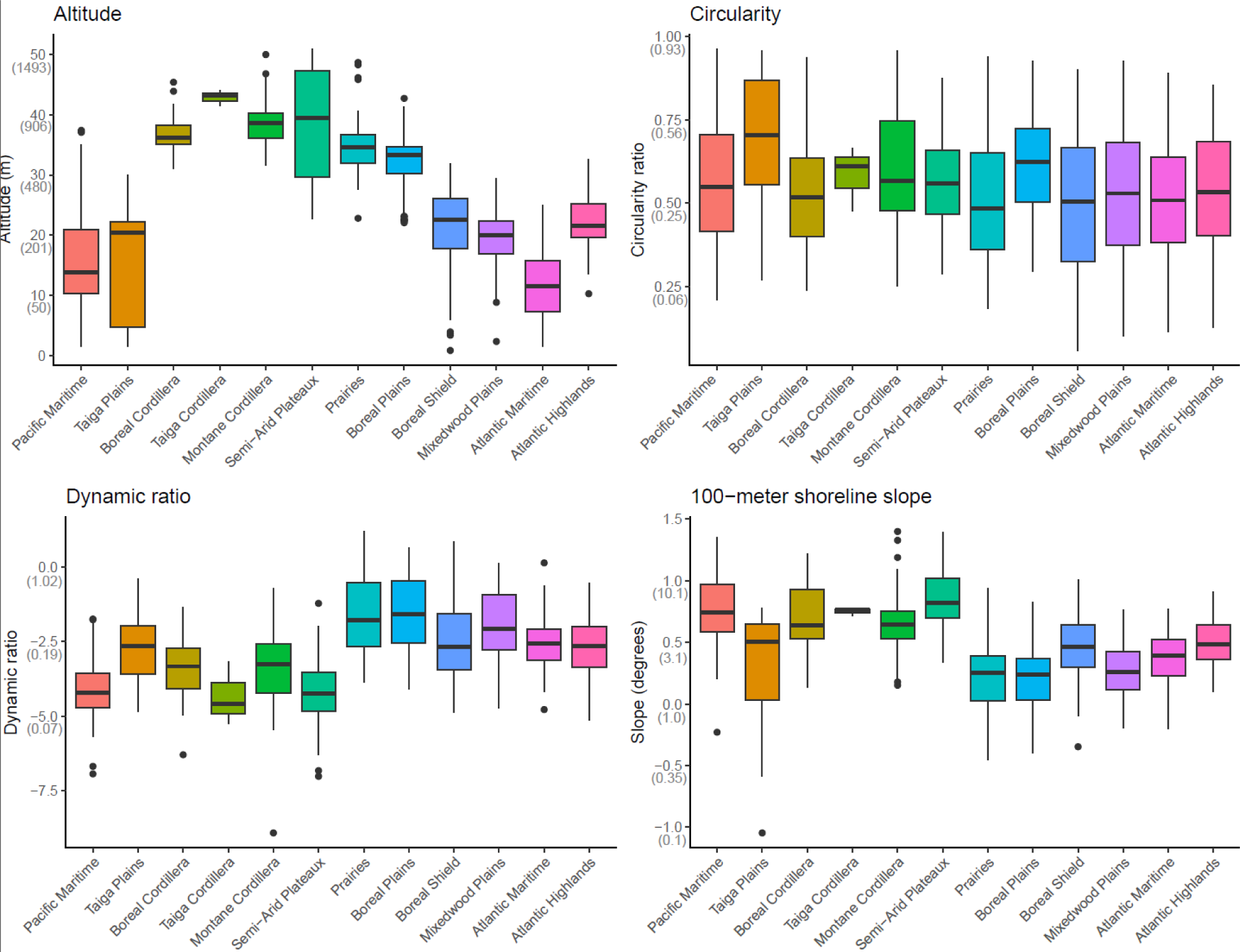
Supplemental Figure S2.

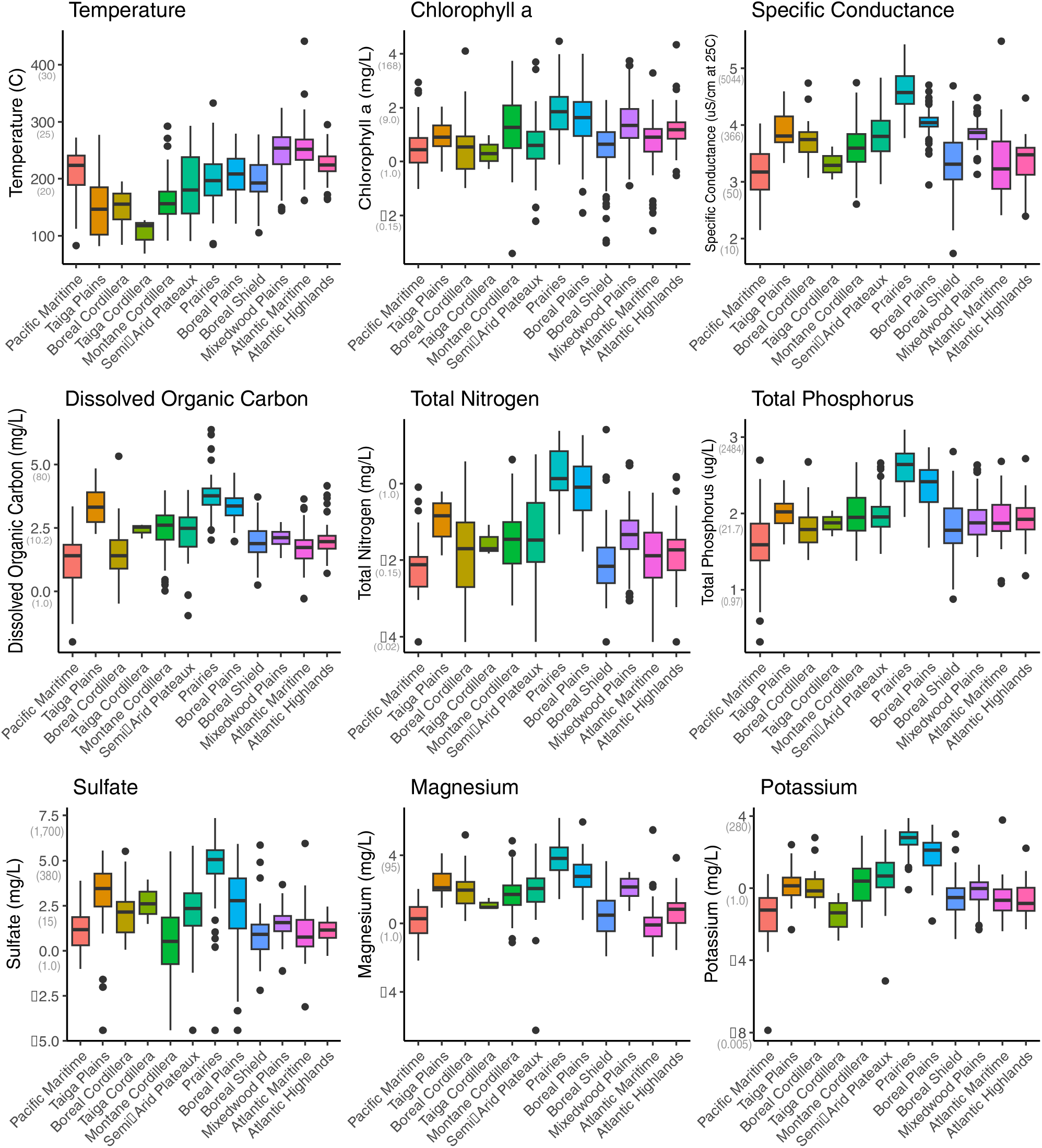
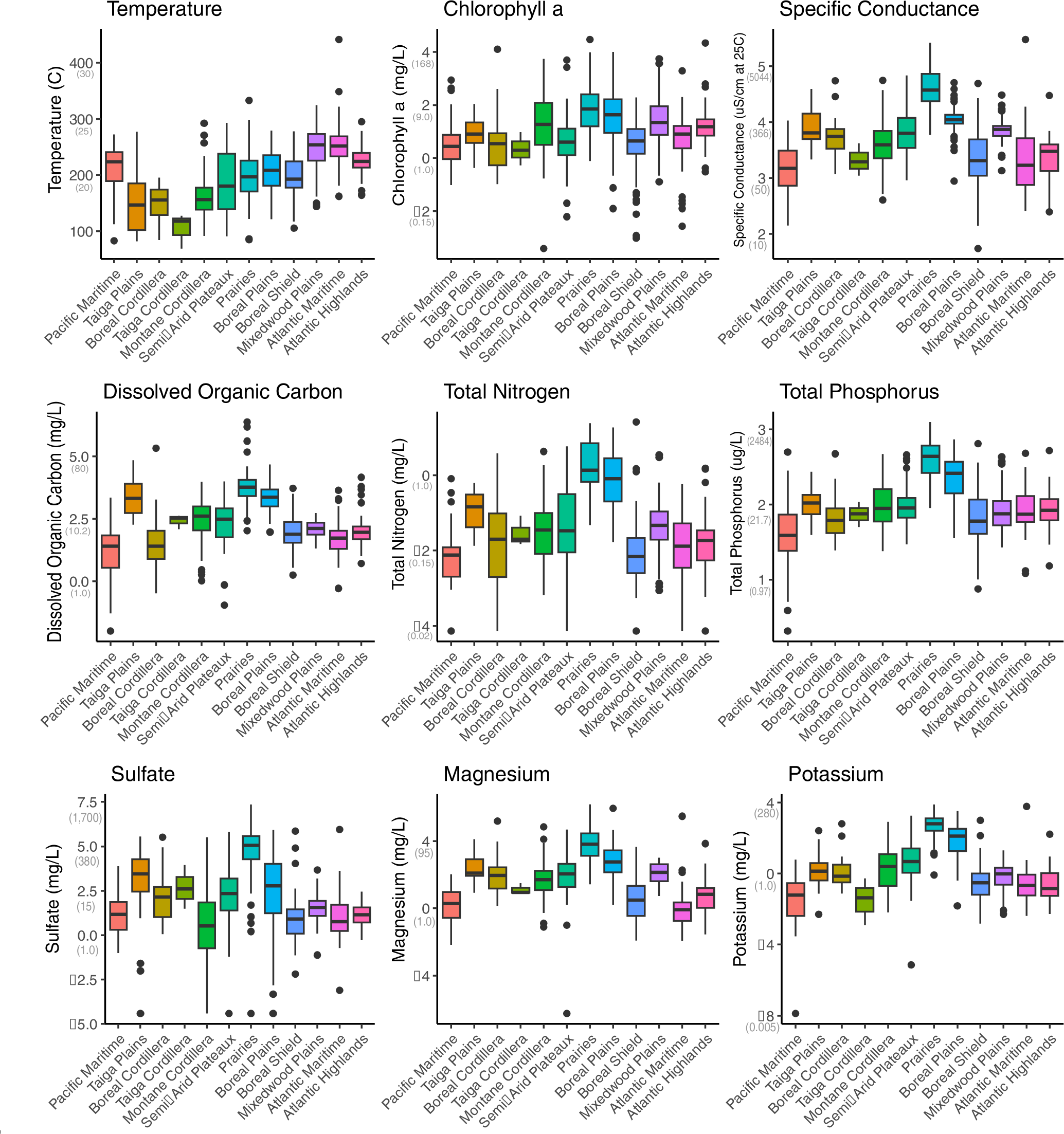
Supplemental Figure S3.

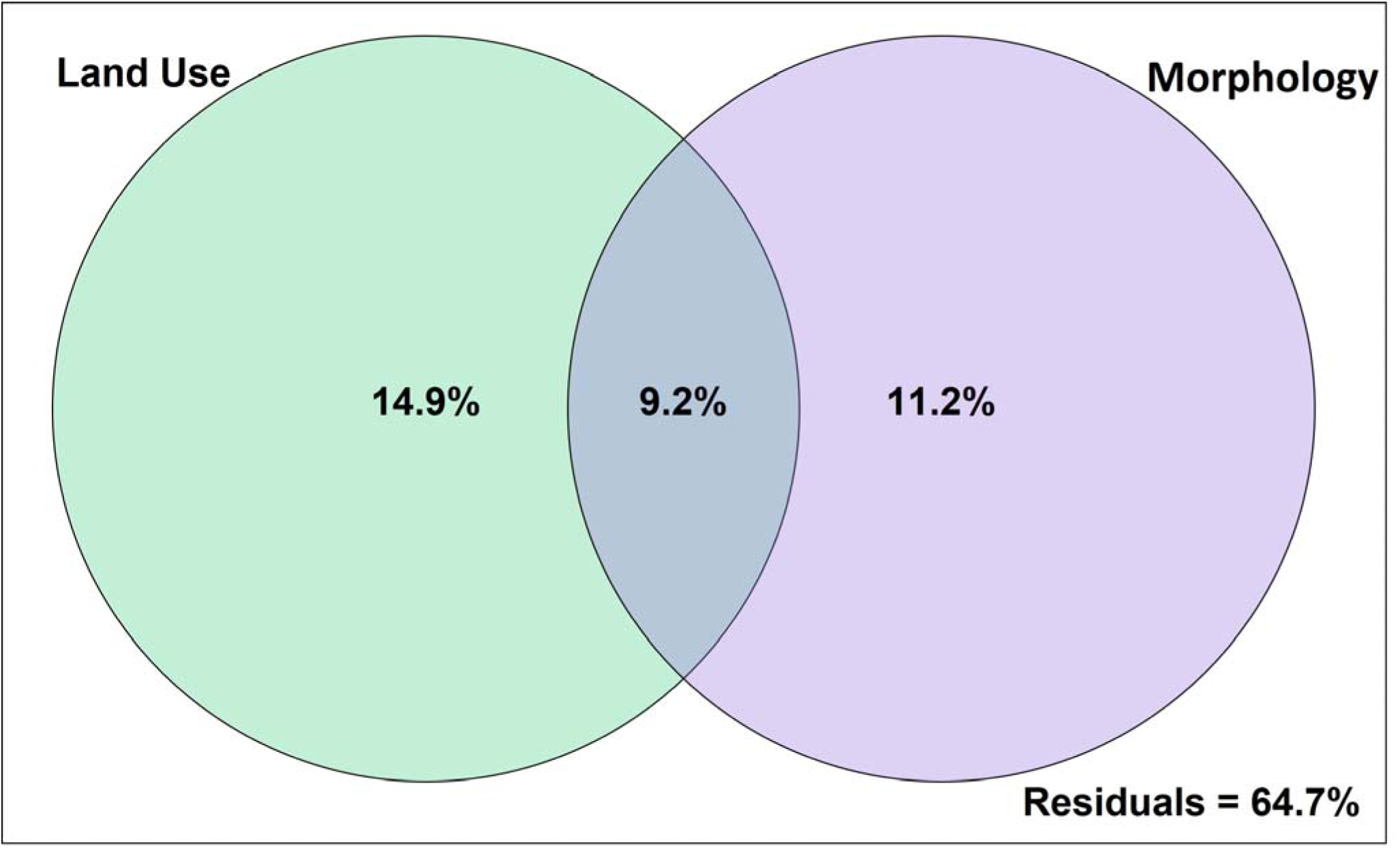
Supplemental Figure S4.

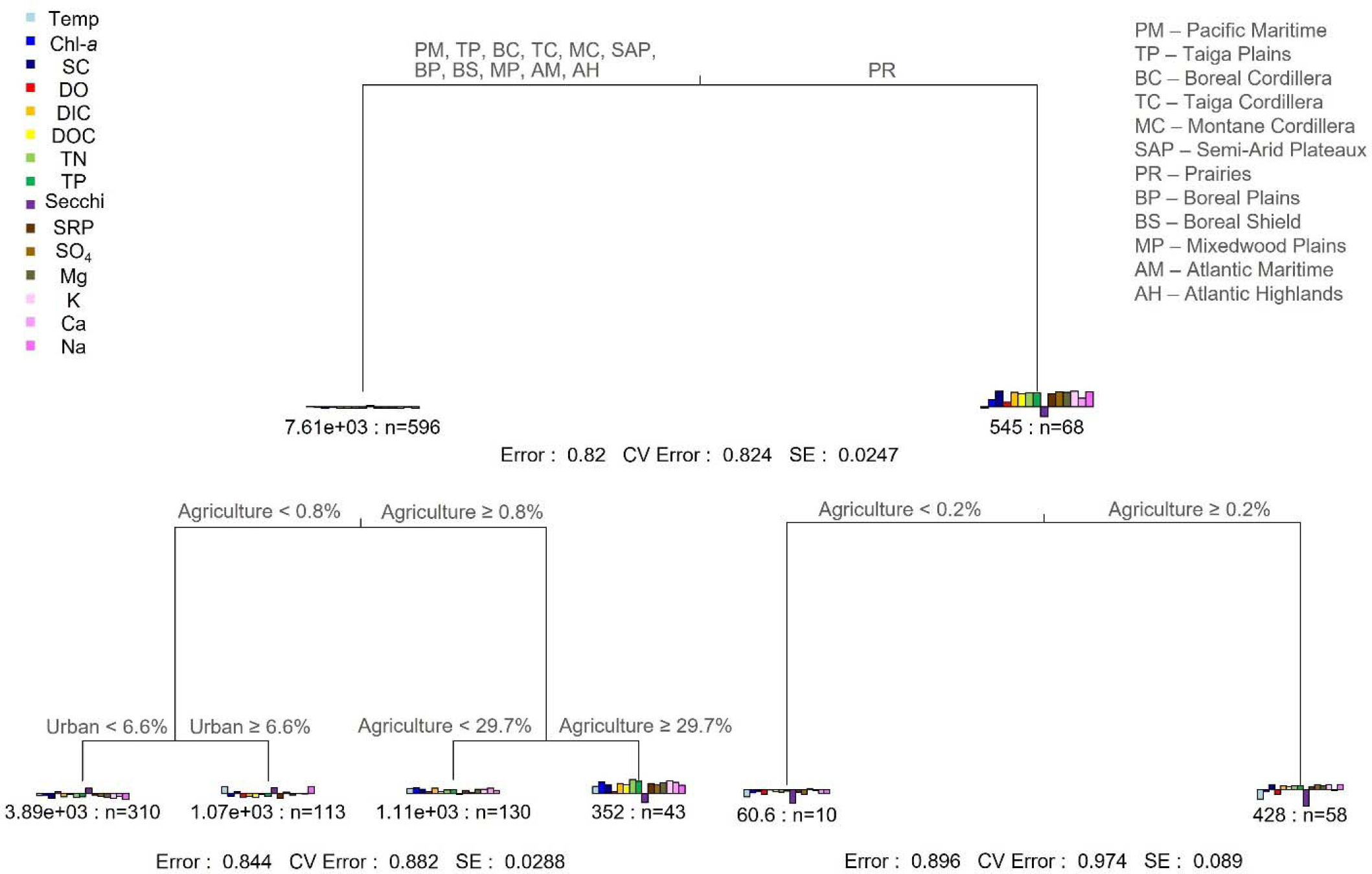
Supplemental Figure S5.

